# CryoROLE: describing large inter-domain rotation in single particle cryo-EM

**DOI:** 10.64898/2026.07.04.736454

**Authors:** Chengmin Li, Wooyoung Choi, Hao Wu, Yifan Cheng

**Affiliations:** Department of Biochemistry and Biophysics, University of California San Francisco; Howard Hughes Medical Institute, University of California San Francisco

## Abstract

In single particle cryo-EM, analysis of continuous conformational heterogeneity has always been challenging. Both linear and deep learning-based methods treat conformational heterogeneity as perturbations to the consensus average conformation, limiting their capability in analyzing large protein motions. While classic conformational classifications are capable of handling large domain motion, they bin continuous protein dynamics into discrete static substates. Here, we present cryoROLE, a computational tool that extracts the continuous conformational dynamics embedded in the static composite map constructed from multi-body refinement into a landscape of relative orientation between the moving domains. Depicted in real space, the landscape allows intuitive interpretations of domain motion and the population of poses in the conformational space. Applying it to various biological systems reveals hidden conformational dynamics that are relevant to protein functions.

## Introduction

With steady technological developments, single particle cryogenic electron microscopy (cryo-EM) has succeeded in achieving atomic resolution routinely with conformationally homogeneous samples, exemplified by reconstructing ∼1Å resolution structures of apoferritin^1,2^. However, many proteins possess various range of local and global conformational dynamics, causing conformational heterogeneity in sample that, if undealt with, deteriorates local or global resolution of the reconstruction. In general, conformational dynamics are continuous, and can be categorized into intertwined two types, deformation (minor or major) and “rigid-body”-like domain motion. The latter is often seen in large macromolecules or complexes comprising multiple domains. Extensive particle classification deals the problem by binning continuous heterogeneity into discrete conformational substates, which can resolve distinct major substates but ignoring subtler heterogeneity within these substates. When a reasonable consensus averaged structure can be generated, continuous heterogeneity (or deformation) can be treated as perturbations to the consensus structure, as demonstrated by both linear hyperspace approach and deep generative latent space methods^3-7^. A classic example is the ratchet motion between small and large subunits in ribosome during active translation^8-11^, which is extensively studied by many methods.

With very large domain motion, such as seen in human fatty acid synthase (hFASN), where the condensing and modifying wings are connected by a short linker and the two wings rotate extensively around each other^12,13^, no reasonable consensus average reconstructions can resolve both domains at good resolution simultaneously. Instead, refining each wing independently by a focused procedure with the other wing being masked out or removed computationally, a composite map can be produced by placing separately refined domains in high resolution together (Fig. 1a, b)^14^. Nowadays, generating a composite map from focused refinement is a commonly used practice in single particle cryo-EM^15^. However, information about relative motion between these domains, which could be functional significant, is lost in such composite maps.

**Figure 1.**
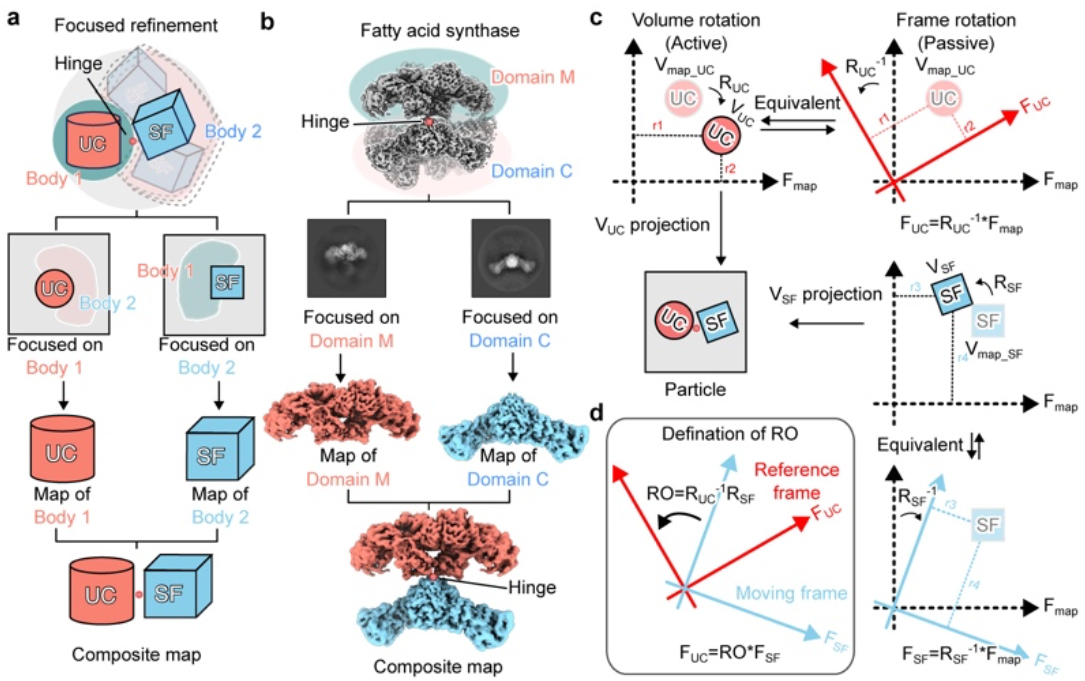
Definition of relative orientation. **a**, Standard focused refinement scheme to generate a composite map. “UC-SF” depicts a two-body object with two domains (UC and SF) linked by a hinge with substantial inter-domain motion. Independent focused refinements on UC and SF yield high-resolution maps for each domain. Combining them produces a composite map with optimal resolution for both bodies. **b**, A corresponding FASN example illustrates the same workflow on experimental data. Domain M and Domain C denote the modifying and condensing wings of FASN, respectively. **c**, Description of active verses passive domain rotation. The same physical rotation can be described either actively by rotating the object in a fixed coordinate frame, or passively by rotating the coordinate frame with object fixed. These two descriptions are mathematically equivalent but using inverse rotation matrices. In the passive rotation, focused refinement assigns each domain a local frame relative to the map frame. **d**, Under the passive rotation, the RO between the two domains is defined as an operation that rotates the coordinate frame of the moving object to the frame of the reference object.

Here, we introduce cryoROLE, a tool to map and visualize particle distributions of relative orientations between two or more domains within a macromolecule. Once a composite map is determined from the same particle dataset, each particle has more than one set of Euler angles, one assigned to each individually refined domain. Thus, for each and every particle, we compute the relative orientation (RO) between these independently refined domains from the Euler angles assigned to them. Plotting RO of all these particles in a Cartesian coordinate system produces a particle distribution landscape that can be visualized and interpreted directly. Such a landscape reveals the range of inter-domain motion and the population of particles at any specific orientation. It extracts conformational dynamics embedded in composite maps and links the substates with continuous domain motions. In practice, cryoROLE enables uncovering protein dynamics that are functionally important but not captured by traditional classification, multibody refinement or deep learning based heterogeneous refinement procedures.

## Results

### Relative orientation between two “rigid bodies”

As defined by the Central Section Theorem^16^, when calculating a three-dimensional (3D) reconstruction in cryo-EM, the Fourier transform of a particle image is inserted into a 3D volume in reciprocal space by a rotational operation *R* corresponding to a specific orientation defined by three Euler angles, ϕ, θ, and ψ, following the convention defined slightly differently in different program packages, such as ZYZ in RELION^17^. Mathematically, particle rotation can be carried out by two equivalent operations: One is the active rotation, in which the coordinate frame, or simply “frame”, is fixed and the particle is rotated by the rotation matrix *R*. The other is the passive rotation, in which the particle is fixed but the frame is rotated by the inverse rotation matrix *R*^-1^ (Fig. 1c). The passive rotation is commonly used in cryo-EM image processing, such as in RELION and cisTEM^18^. Here, we treat the orientation metadata as the passive rotations.

In a two-body refinement procedure (Fig. 1a, b), each domain of a particle is assigned with a set of Euler angles by refining it against its corresponding volume within the composite map. Thus, each particle in the dataset that produces the composite map has two sets of Euler angle assignment. Choosing one domain as the reference and the other as the moving domain, the relative orientation (RO) between the two domains in a specific particle is calculated as the matrix product (Fig. 1d):

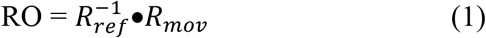

The RO defined here is invariant to global reorientation applied equally to both domains, such as re-orienting the composite map, or micrograph-based alignment offsets, such as residual beam tilt, imperfect motion correction or CTF model. Consequently, RO reflects only the true inter-domain orientation relationship, along with any domain-specific uncertainties generated during refinement, such as alignment errors or one domain is less well resolved than the other, etc.. Thus, RO describes per-particle inter-domain orientation when both domains behave as “rigid bodies”. Detailed descriptions are in Methods.

### RO landscape and cryoROLE

Calculating RO of all particles used to generate a composite map and plotting the result in a Cartesian coordinate system produces a particle distribution landscape of RO between two “rigid” domains (Fig. 2). We implemented the procedure of calculating and displaying RO landscape into an open-source Python package, cryoROLE, stands for <u>cryo</u>-EM <u>R</u>elative <u>O</u>rientation <u>L</u>andscap<u>E</u>. With two per-domain metadata (.*star* of RELION or .*cs* of CryoSPARC), CryoROLE computes per-particle RO, pools the results into a landscape in the rotation vector (RV) space for easy statistical analysis, and displays it in a fixed-axis Euler-angle coordinate system, which we refer to Euler-angle space, for visualization and intuitive interpretation. In this coordinate system, α, β and γ correspond to sequential rotations around the fixed Z, Y, and X axes, respectively, that define the RO between the two domains (Fig. 2b). Optionally, cryoROLE canonicalizes the coordinate frame by aligning α to represent the primary rotation and β as the secondary rotation, respectively. We refer this procedure as frame canonicalization and the resulting coordinate system as canonicalized frame. The landscape is color coded with the local particle density estimated from scaled local density (SLD) in RV space.

**Figure 2.**
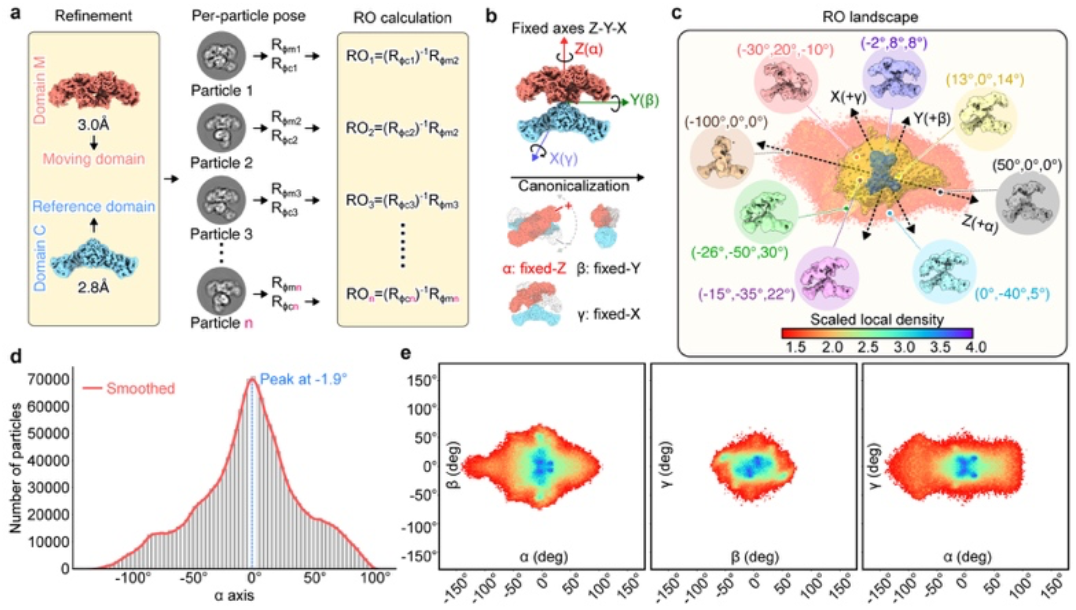
RO landscape by cryoROLE. **a**, Focused refinements of domain C (condensing wing, as reference domain) and M (modifying wing, as moving domain) of FASN yield two orientation assignments, one for each domain of every particle, from which RO between two domains is calculated for all particles in the dataset. **b**, All ROs are expressed into fixed-axes Z–Y–X Euler angle space as α, β, γ for visualization. After frame canonicalization, rotations around the fixed Z, Y, and X axes correspond to the primary, secondary, and tertiary motion directions, respectively. **c**, RO landscape of FASN is viewed in 3D. Each point represents one particle, and the color indicates scaled local density (SLD) estimated in rotation-vector space. Panels are displayed after SLD filtered with a threshold of 1.3. Eight representative coordinates (α, β, γ) were selected from the landscape, and particles within a 7°–9° geodesic radius were extracted for reconstruction without further refinement. **d**, 1D profile of particle number distribution, both histogram (bar) and kernel-density estimate (KDE, smooth red curve), along the primary motion coordinate a. **e**, 2D projections of the RO landscape onto the α–β, β–γ, and α–γ planes.

Depicted in a Cartesian coordinate system of a physically interpretable real space, the RO landscape allows for intuitive direct interpretation, which is different from the hyperspace projection of principle component analysis (PCA) or the latent space depicted in deep learning based heterogeneous refinement programs. Coordinate of each and every point in RO landscape defines a particle with a specific orientation between two domains. Points located near each other in the landscape, as evaluated by geodesic distance in RV space, represent particles with similar conformation between the two domains. The number of particles in a local region of the landscape represent a relative population of specific RO in the entire dataset. Therefore, cryoROLE converts per-particle poses into a global description of how two domains move with respect to one another across the dataset. Together, the landscape generated in this way describes the dynamic behavior of the complex in terms of the orientation variations between its two domains.

### Validation of cryoROLE

CryoROLE was initially developed to analyze the dynamics behavior of hFASN, which contains two large wings rotate around each other^13^. With large rotation between two wings, other methods, such as conventional extensive particle classification, 3D variability analysis^4^ or cryoDRGN^7^, failed to describe the dynamic behavior of hFASN. Because both wings are recognizable in a reconstruction, even at the modest resolution, hFASN dataset is ideal for validating the formulation used in cryoROLE and demonstrating various features of the landscape. For this purpose, we combined all hFASN datasets collected with different conditions into one with ∼2.2 million particles, reprocessed it with RELION, and calculated RO landscape by cryoROLE (Fig. 2a, Supplementary Fig. 1a).

A key validation of the landscape is that particles selected from any specific location in the landscape should produce a reconstruction that, without any further refinement, show the two domains in a relative orientation defined by the coordinate in the landscape. Indeed, selecting particles from any position in this landscape produce a reconstruction that has the RO between two wings as defined by that coordinate (Fig. 2b, c). Projecting the 3D landscape into a 2D plane or further into a 1D profile makes it easier to visually interpret the landscape (Fig. 2d, e)

Using hFASN dataset, we tested various features of the landscape. First, exchanging reference and moving domains in the composite map results in flipping the landscape without any other changes (Supplementary Fig. 2a, b). The shape and overall particle distribution profile in RO landscape is not impacted by the total number of particles, as far as they are sufficient to achieve a reasonable resolution for each domain (Supplementary Fig. 2c, d). Thus, landscapes obtained from different datasets of the same sample obtained under different experimental conditions can be compared directly. Second, by placing the axis of α rotation along the primary rotation, the canonicalized RO landscape is fully defined regardless the orientation of the composite map (Supplementary Fig. 3). Third, resolution of the reconstruction calculated from extracted particles within a defined geodesic radius from a specific coordinate depends on particle numbers being selected and the geodesic distances used to extract the particles. Extracting particles from a smaller geodesic radius yield fewer particles to calculate the reconstruction, while from a larger region provides more particles but with increased conformational heterogeneity (Supplementary Fig. 4a). Also, particles selected from some coordinates in the landscape may show impossible domain configuration, such as overlapping of two domains, likely caused by mis-alignment (Supplementary Fig. 4b).

In the following, we apply cryoROLE in two different biological systems, to extract information that are otherwise buried, and to demonstrate various features of the landscape.

### Chromatin remodeler INO80 with hexasome

ATP dependent chromatin remodeler INO80 slides its substrates, either nucleosome or hexasome, along DNA. In our previous cryo-EM study of the INO80-hexasome complex^19^, we were able to refine INO80 to 2.8Å resolution, but with poor hexasome density. Subsequent 3D classification focused on hexasome identified three conformational classes. For each one, separate refinement of INO80 and hexasome produced three composite maps, corresponding to placing Ino80 ATPase domain at near superhelical location (SHL) -2, -3 and in between, -2.5. Here, we combined particles from these three classes and reanalyzed the data. We refined INO80 and hexasome individually to nominal resolution of 2.8Å and 3.1Å, assembled a single composite map (Fig. 3a). Subjecting the per-particle orientation metadata from the two focused refinements to cryoROLE produced a RO landscape. After motion axes canonicalization, the α axis describes the dominant rotation of INO80 around the hexasome. This landscape shows that particles are continuously distributed along α in an elongated region spanning approximately 110° but narrow in the other orthogonal directions (Fig. 3b, Supplementary Fig. 5). The corresponding position of three classes identified from original classification are mapped onto the landscape. One-dimensional (1D) profile along α shows several peaks (Fig. 3c), revealing a continuous trajectory with preferred orientations rather than a uniform distribution.

**Figure 3.**
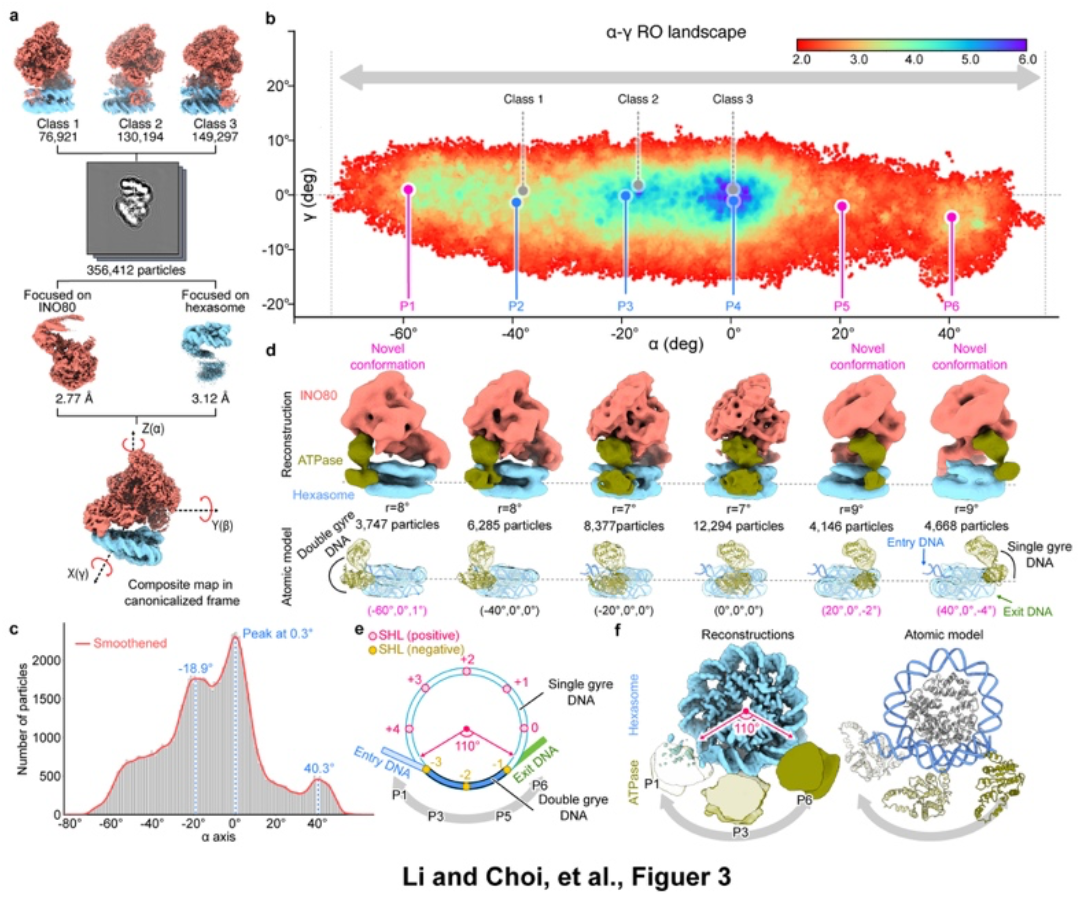
RO landscape of INO80–hexasome complex. **a**, Particles from three previously classified INO80-hexasome states were merged and reprocessed with focused refinements and yielded an initial composite map. The composite map was reoriented into the canonicalized coordinate frame in cryoROLE, in which rotations around the fixed Z, Y and X axes correspond to α, β and γ, respectively are marked. **b**, Canonicalized RO landscape of INO80-hexasome displayed as a 2D projection in the α-γ plane of fixed-axis Z-Y-X Euler-angle space. The positions of the three input classes and six representative landscape coordinates along the α axis, from -60º to 40º at 20° intervals, P1–P6, are indicated. The panels are displayed after SLD filtered with a threshold of 2.0. **c**, 1D profile of particle number distribution, both histogram (bar) and kernel-density estimate (KDE, smooth curve), along the primary rotation axis, α. Local peaks are located at -18.9°, 0.3°, and 40.3°, corresponding to placing Ino80 ATPase at near SHL -2.5, -2 and -1. The positions of the three input classes are indicated. **d**, Reconstructions calculated using particles extracted within a 7– 9° geodesic radius around P1 to P6. All reconstructions are displayed with hexasome in the same orientation. Ino80 ATPase domain can be localized. INO80 density is shown in salmon, the hexasome in sky blue, and the ATPase lobes in olive. From P1 through P6, the ATPase density progressively shifts from the entry-DNA side to dyad of the hexasome. **e**, A schematic diagram of hexasome showing the positions of P1-P6 on the hexasome. **f**, Densities of hexasome with Ino80 ATPase at P1, P3 and P6 position (left) and the corresponding atomic models (right).

To visualize how INO80 rotates on the hexasome, we sampled the RO landscape along α rotation from −60° to 40° in 20° increments, P1 to P6. For each position, we selected particles within a 7° ∼ 9° geodesic radius around the corresponding landscape coordinate and reconstruct without further refinement. All six reconstructions show well-resolved density for both INO80 and the hexasome, enabling direct fit with PDB structures (Fig. 3b, d). These reconstructions correspond to placing Ino80 ATPase at different position along hexasomal DNA (Fig. 3e, f). P2, P3 and P4 correspond to Ino80 ATPase near SHL-3, -2.5 and -2 position. However, in P6, the Ino80 ATPase is positioned near SHL -1, which is a low-occupancy state that was not identified as an independent class in the original classification. With the continuous particles distribution along α axis, the landscape provides a trajectory of how the remodeler INO80 could rotate around hexasome. Across P1 to P6, the Ino80 ATPase relocates continuously from the entry-DNA side toward dyad, consistent with the INO80 remodeling model that was proposed^19^.

The DNA segment connecting the entry and exit sides contains only “double-gyre” DNA region on the hexasome, spanning approximately from SHL −3 to slightly beyond −1 positions (∼110°). Interesting, this segment coincides with the full range of particle distribution along the α in the landscape (Fig. 3e). Because these data were collected in the absence of ATP, this result suggests that, without ATP-driven remodeling, the ATPase remains confined to this double-gyre DNA segment.

### Ribosome after thermo-annealing

During protein synthesis, ribosome undergoes intersubunit rotation between small (SSU) and large (LSU) subunits which is often referred to as ratchet-like rotation^9^, together with swiveling of the head of SSU relative to the SSU body during tRNA-mRNA translocation^20^. These motions are depicted by the ribosome angle decomposition^10^. Without being sorted out either computationally or biochemically, such conformational heterogeneities deteriorate reconstruction quality. In an earlier study of factor-free apo 70*S* ribosome from *Escherichia coli*, it was found that temperature annealing prior to cryo-EM grids preparation homogenizes ribosome conformational states, improving the quality of reconstructions^21^. Here, we applied cryoROLE to this published dataset to derive landscapes of rotational motion between different domains of ribosome, and to reveal how temperature annealing changes the landscapes of such inter-domain motion.

We obtained two datasets from the authors of Chu et al.^21^, one is of *E. coli* ribosome collected without temperature annealing (655,257 particles) and the other after temperature annealing (509,941 particles). Combining them together, we re-processed the data by a common workflow in cryoSPARC with focused refinement on three ribosomal bodies: the LSU, SSU-body, and SSU-head, reaching nominal resolutions of 2.28Å, 2.44Å and 2.56Å, respectively, generating a single composite map (Fig. 4a). Instead of using the default canonicalized axes, we oriented the composite map defined by ribosome angle decomposition^10^ (Fig. 4b). In the resulting RO landscape, the α and β coordinates therefore primarily report ratchet-like and tilt-like components of motion, respectively. Treating the whole particle as three-pairs of two-bodies, we applied cryoROLE to generate RO landscapes between: SSU-body to LSU (B2L, LSU as reference), SSU-head to SSU-body (H2B, SSU-body as reference) and SSU-head to LSU (H2L, LSU as reference) with two datasets combined, as well as decomposed into without and with temperature annealing (Fig. 4c, Supplementary Fig. 6). Processing combined dataset through the same refinement workflow minimized pipeline-dependent differences in pose estimation, making the landscapes of different samples directly comparable to reveal changes of both the ratchet and head swiveling motions within ribosomes in response to temperature annealing. Without annealing, the landscape shows apo ribosome has broader ratchet motion (B2L) and head swiveling (H2B), which after temperature annealing are significantly reduced, as shown in confined hot spots in landscape after annealing (Figure 4c, Supplementary Fig. 6).

**Figure 4.**
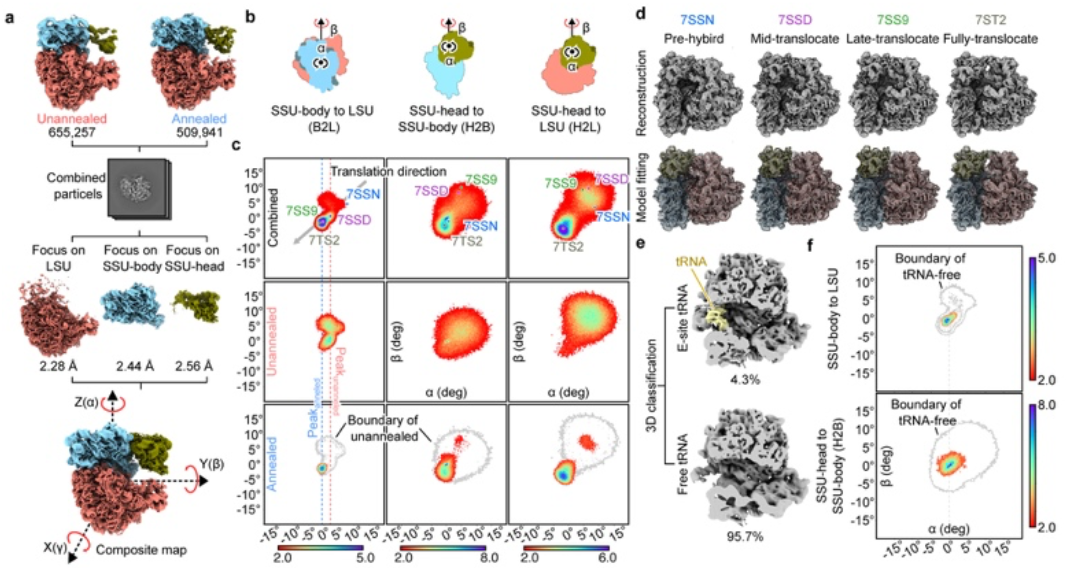
RO landscape of ribosome. **a**, Focused refinement of combined datasets of temperature annealed (right) and unannealed (left) ribosome yielded a composite map. The composite map was oriented according to ribosome angle decomposition, with the fixed Z axis aligned to the canonical inter-subunit ratchet axis and the fixed Y axis aligned to the tilt axis. Rotations about fixed Z, Y and X are denoted as α, β and γ respectively. **b**, Definition of three RO pairs and motion axes: SSU-body to LSU (B2L), SSU-head to SSU-body (H2B), and SSU-head to LSU (H2L). **c**, 2D projections of the RO landscapes for B2L, H2B, and H2L in α-β plane for combined (top), unannealed (middle), and annealed (bottom) datasets. All panels are displayed after SLD filtered with a threshold of 2.0. The annealed sample displays contracted RO distributions and shifts the density peak compared with the unannealed sample. In the combined landscapes, the positions corresponding to four representatives published 70S structures are indicated. **d**, Particles within a 3° geodesic radius of the landscape coordinates mapped from four published 70S structures, 7SSN, 7SSD, 7SS9 and 7ST2, were extracted and reconstructed. Top: reconstructed maps. Bottom: the same maps fitted with the corresponding atomic models. **e**, Focused classification separates the combined dataset into an E-site tRNA-bound subset and a tRNA-free subset. **f**, 2D projections of the RO landscapes of each subset are shown with the dashed outline marks the boundary of the tRNA-free landscape. The E-site tRNA–bound particles occupy a narrower region within the broader tRNA-free landscape, suggesting that E-site tRNA restricts inter-domain motion of the 70S ribosome.

To further assess the interpretability and the functional relevance of the RO landscape, we projected four previously published, translocation-related 70S states onto the combined RO map: pre-hybrid state with A/P* + P/E tRNA before EF-G bound (PDB: 7SSN), mid-translocation state with ap/P + pe/E tRNA after EF-G bound (PDB: 7SSD), the late intermediate (PDB: 7SS9) and fully translocated state with P/P tRNA after EF-G dissociated (PDB: 7ST2) (Figure 4c, combined particle landscape). Once mapped onto the landscape, all these structures fall approximately along a trajectory in B2L landscape, consistent with a one-dimensional progression of inter-subunit rotation during translocation. Furthermore, 7SSD, 7ST2, and 7SS9 are located in the landscape with higher particle populations than that of the pre-hybrid state 7SSN. In particular, the fully translocated state (7ST2) is mapped close to a hotspot, consistent with the idea that the apo dataset after annealing is enriched for a non-rotated, post-translocation-like geometry, reinforcing the idea that RO landscape represents the thermodynamic preference of the system during its functional cycle (Fig. 4c). Thus, known functional states can be used to anchor the RO landscape and facilitate mechanistic interpret with the experimental particle distribution (Supplementary Fig. 7).

Around each mapped coordinate, we selected particles within a 3° geodesic radius and generated reconstructions without further refinement. In all four cases, the resulting maps have well resolved LSU, SSU-body, and SSU-head, with the corresponding deposited atomic models fit well into the maps (Fig. 4d).

From the combined dataset, we also performed a conventional focused 3D classification to separate particles into an E-site tRNA-bound population (4.3%) and a tRNA-free population (95.7%). We then computed RO landscapes for each population. In both B2L and H2B landscape, the E-site tRNA-bound particles are concentrated at the core region of the landscape, whereas the tRNA-free particles exhibit a broader distributed. This indicates that, while the ribosome possesses intrinsic flexibility, the presence of tRNA restricts the global motion of both the SSU head and body, locking the ribosome into a narrower range of orientations (Fig. 4e, Supplementary Fig. 8).

## Discussion

Protein dynamics, which are often tightly associated with protein function, is one of major sources of conformational heterogeneity encountered in single particle cryo-EM. In recent years, computational heterogeneous analysis has become a major theme of research in cryo-EM. Recent examples include principal component analysis (PCA) based 3D variability analysis (3DVA)^4^, deep learning based cryoDRGN^7^ and DRGN-AI^3^, physics-informed deep learning method 3DFlex^5^, and Gaussian pseudo-atomic model based DynaMight^6^, etc.. These methods differ substantially in their model assumptions and output representations, but they share the aim to describe conformational heterogeneity caused by flexible motion in macromolecules. They can also be applied to deal with “rigid-body”-like motion, such as ratchet motion in ribosome. However, when applied to hFASN and INO80-hexosome, where conformational heterogeneities are no longer caused by small perturbation around a consensus conformation but dominated by large “rigid-body” like motion between domains, available continuous-heterogeneity methods are unable to produce satisfactory results. Instead, classic approach of intensive classification followed by focused refinement and composite map generation is more robust and practical. But it bins the heterogeneity into discrete conformational substates rather than continuum. CryoROLE extends this classic workflow by converting the per-particle poses from focused refinements into a continuous RO landscape to describe very large and continuous domain motion of a protein complex, either intrinsic motion or driven by biological processes. Computationally, it is a pure mathematics-based algorithm without pre-training and requires minimal computational resources.

The closest related procedure of cryoROLE is the “multi-body refinement” implemented in RELION^14^, which defines a set of independently moving rigid bodies and refines the pose of each one by combining focused refinement with iteratively updated partial signal subtraction. After convergence, the RO between these bodies are further analyzed by PCA. Thus, both multi-body refinement and cryoROLE extract the RO information from volumetric data instead of projection images. The difference is that cryoROLE avoids PCA but maps the full distribution of RO of each particle directly so that the landscape is intuitively interpretable. In practice, cryoROLE takes the .star files from multi-body refinement to generate the landscape. It can also take the results from other focused refinement procedures, either traditional manual analysis or automated pipelines such as cryoSPARC. For both hFASN and INO80-hexasome complex, where the automated multi-body refinement procedure failed, the composite maps were generated by performing traditional focused refinements of individual domains, from which cryoROLE produced landscapes.

Because the landscape generated by cryoROLE is in real space, it is straightforward to compare landscapes of the same complex under different biochemical or physical conditions, which is less straightforward in latent or eigenvector spaces. Considering that the distribution of particle population in the landscape is related to the relative preference of a specific orientation, arguable, it is related to the thermodynamic free-energy landscape. CryoROLE requires reliable per-domain pose estimates from focused refinements of rigid body domains. Arguably, no domain motion during any biological processes is rigid-body motion. Also, it does not deal with compositional heterogeneity. These limitations can be mitigated by combining cryoROLE with other heterogeneous analysis methods. For example, large motion can often be described as combinations of rigid-body motions and local domain deformations. Thus, a plausible more comprehensive approach is to generate a local consensus reconstruction with particles selected from confined area in the RO landscape, followed by treating the flexible motion as perturbations to this local consensus map (Supplementary Fig. 9). Similarly, cryoROLE can be applied after 3D classification to isolate compositional heterogeneity, as demonstrated here (Fig. 4e).

## Methods

### Description of cryoROLE

#### Description of relative orientation (RO)

Considering a macromolecule that contains two near-rigid domains, one is chosen as the reference domain and the other as the moving one. For particle *i*, focused refinements of these two domains provide two sets of parameters, either Euler angles as in RELION^17^ and cisTEM^18^, or rotation vectors as in cryoSPARC^22^, that describe the orientation relationship between the projection image and the reference volumes in the composite map. With each parameter set, a rotational matrix, *R*_ref_(*i*) for the reference domain and *R*_mov_(*i*) for the moving domain, is expressed under the same composite-map coordinate frame, ℱ. Throughout this work, we use the passive, frame-based interpretation of the rotational matrices. Under this convention, new frame of each domain matching the projection to the reference volume in the composite map are obtained as:

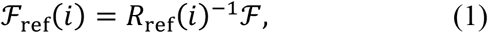

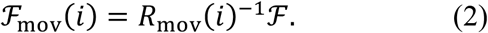

Thus,

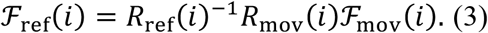

The relative orientation (RO) between the moving and reference domain is defined as a rotational operation that matches the moving-domain frame into the reference-domain frame:

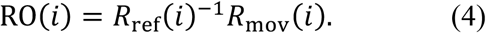

In a 3D space, the orientation of each domain has three rotational degrees of freedom. Thus, a two-domain system contains six degrees of freedom. Because two domains are constraint in one particle, it cancels out three rotational freedoms, leaving three degrees of freedom represented by RO(*i*) ∈ *SO*(3). RO is therefore a minimal descriptor of the relative rotation between two near-rigid domains within a particle.

Conceptually, RO is the per-particle inter-domain rotational relationship, which is intrinsic to the particle and is invariant of any global reorientation *G* applied equally to both domains:

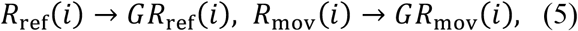

and

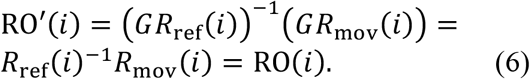

Thus, once a reference composite map is defined, RO between the moving and reference domain of every particle in the dataset is defined. Same, exchanging the reference and moving domains inverts but does not change RO:

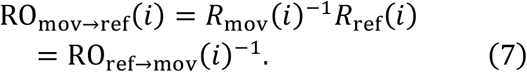

#### Description of RO landscape

A landscape is generated by plotting RO of all particles that generate the composite map as point clouds in a 3D space. To do so, RO needs to be numerically represented as a 3×3 rotation matrix, a rotation vector or a set of Euler angles, and expressed in a chosen coordinate frame. Being able to express RO in different format and to convert between them allow various robust computational operations. For example, rotation vectors provide convenient coordinates for local statistical analysis and frame canonicalization, and Euler angles for visualization and interpretation.

Same as reorienting the reference composite map, reorienting the coordinate frame does not alter RO in the particle either. Changing the coordinate frame by a rotation C:

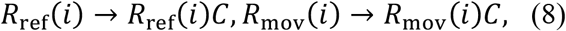

Then the same physical RO is expressed as its conjugate:

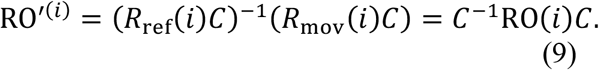

Thus, regardless the choice of the coordinate frame of the RO landscape, its intrinsic geometry and properties are preserved, including pairwise rotation distances, local neighborhoods, density relationships and cluster structure. This is important for comparing datasets or processing workflows that start from differently initial composite map, which may produce different coordinates for the same physical RO (Supplementary Fig. 3a). This also allows cryoROLE to perform frame canonicalization (Supplementary Fig. 3b).

#### Statistical analysis in rotation vector (RV) space

A rotation vector (RV) is the axis-angle representation of a rotation, defined as the product of a unit rotation axis and the corresponding rotation angle. For each particle, we convert the RO matrix RO(*i*) into a rotation vector

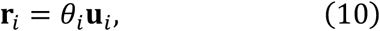

where **u**_*i*_ is the unit rotation axis and *θ*_*i*_ is the rotation angle in radians. The Cartesian components of **r**_*i*_ are denoted as (*r*_*x*_, *r*_*y*_, *r*_*z*_) . Unlike Euler-angle coordinates, RV coordinates are locally Euclidean and do not suffer from Euler-angle gimbal singularities. We therefore perform statistical analysis, coordinate re-mapping and neighborhood-based operations in RV space, while using Euler-angle coordinates primarily for visualization and interpretation.

#### Canonicalization of coordinate frame

Because all intrinsic properties of a landscape are preserved regardless the choice of its coordinate frame, we can canonicalize the coordinate frame by rotating Z axis in the rotation vector space to the direction representing the largest variation in the landscape, and Y to the largest variation while being perpendicular to Z. Consequently, changing the coordinate frame rotates the entire RV point cloud without changing its intrinsic geometry. CryoROLE uses this property to canonicalize the landscape by rotating the coordinate frame so that its Z, Y and X axes correspond to the primary, secondary and tertiary directions of variation in the RV point cloud. This canonicalized RO landscape is then represented in an extrinsic, fixed-axis ZYX Euler angles space, such that the dominant rotational variation is represented mainly by α (rotation about fixed Z), followed by β (fixed Y) and γ (fixed X). The same frame transformation is applied to the composite map, allowing the landscape axes to be interpreted directly on the structure. This canonicalization improves interpretability of the landscape and its comparability with that generated from other datasets, being processed by different workflows, or begin from differently oriented composite maps.

#### Scaled local density (SLD)

To quantify local particle density in the RO landscape, cryoROLE computes the nearest-neighbor-based score for every particle in RV space, which is defined as scaled local density (SLD). Let

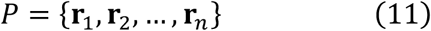

be the set of all per-particle RV coordinates. For point **r**_*i*_, the Euclidean distance *d*_*ij*_ denote its *j*-th nearest neighbor in RV space, computed using a k-d tree. The mean distance of the first *k* nearest neighbors around **r**_*i*_ is:

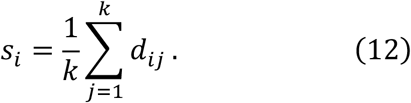

Denser regions have smaller *s*_*i*_, whereas sparse regions have large *s*_*i*_. We further define a global mean distance of nearest neighbor as,

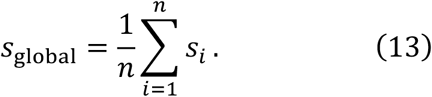

So that the SLD for particle *i* is defined as

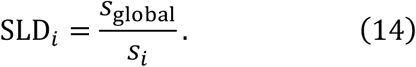

Values greater than 1 indicate that the local particle distribution is denser than the global average, below 1 indicate a sparser than average distribution. In cryoROLE, the default setting is *k* = 30. Using SLD to present the particle population distribution of RO landscape, the distribution profile is not impacted by the total particle numbers, as far as the total number of particles is sufficient to produce well a defined composite map. Thus, landscapes between datasets with different total number of particles used to reconstruct the composite maps can be compared.

#### Landscape visualization in fixed-axis Euler angle space

After coordinate frame canonicalization, local density estimation, each RO is rendered using the fixed axis Z-Y-X Euler angle coordinate expression, and color coded by SLD. Thus, the RO matrix of every particle in the landscape is described as

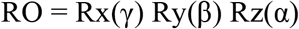

where Rz, Ry and Rx denote angles that rotate the moving domain non-commutatively to its corresponding orientation in the composite map. This display is visually intuitive of how the moving domain is oriented relative to the reference domain.

We rendered the landscape in 3D space, which can be visualized by UCSF ChineraX as point cloud, in which each particle is displayed as a point with SLD as its value. The point cloud can be converted into a volumetric density map in MRC format, with its pixel value as the average SLD of all particles fall within the pixel. Such conversion enables standard map visualization, such as isosurface contouring and orthogonal slicing, etc. The volumetric map can be projected into 2D for presentation or into 1D to show particle population distribution profile along one specific axis.

Particles around a specific coordinate can be extracted for further analysis or volume reconstruction. Particle extractions are performed in RV space, using geodesic angular distance around a specific location in the landscape. Extracting particle in RV space avoids artifacts caused by Euler angle wrapping, parameterization nonlinearity or gimbal singularities. Extracted particles carry two .star files from the input, but a reconstruction calculated with one .star file has two domains oriented as defined by the coordinate of where the particles are extracted. Depends on the geodesic distance used to extract particles, the volume corresponding to one .star file typically have a better resolution than the other volume. Extracted particles can also be subject to heterogenous analysis by other programs (Supplementary Fig. 4a and 9).

### Validation of cryoROLE

CryoROLE was originally developed to describe the rotational motion between two wings of human FASN^13^. We combined all datasets of hFASN from previous study into a single dataset of 2,234,482 particles and reprocess it. After performing focused refinement as described^13^, a RO landscape was produced using the released version of cryoROLE, in which the condensing wing was chosen as the reference domain and the modifying wing as the moving domain. Using this landscape, we validated the formulations used to generate the landscape, and all features described above.

Exchanging the reference and moving domains inverted the RO: in RV space, each rotation vector **r** was mapped to −**r**, while inter-point distances, local support and clustering were preserved. The corresponding RO landscape in EA space shows the same transformation (Supplementary Fig. 2a, b). Applying various global rotations to the composite map resulted in corresponding rotations of the landscapes in the RV space, which after canonicalization are aligned with each other. Fixed-axis Euler angle display coordinates derived from these aligned RV coordinates were therefore directly comparable. After canonicalization, the hFASN landscape showed a spread largely along the α coordinate, whereas the spreads along β and γ were narrower. Particles selected from around various coordinate in the landscape produced reconstructions with the defined orientation between condensing and modifying wings. Furthermore, randomly retaining 10% of the full dataset preserved the overall landscape and produced an SLD-colored distribution similar to that of the full dataset when visualized with the same settings (Supplementary Fig. 2c). Further reduction of particles to a 1% subset still retains the approximate shape and boundary of the landscape but with noticeably less stable SLD values, reflecting insufficient sampling of local neighborhoods (Supplementary Fig. 2d). These tests indicate that SLD is most reliable when the underlying orientation distribution is adequately sampled and when datasets are processed with comparable settings.

To calculate reconstruction from particles selected from the landscape, there is a practical trade-off between angular homogeneity and numbers of the selected particles. We selected particles from a specific fixed-axis Euler coordinate (α, β, γ) = (−2°, 8°, 8°) using geodesic radii from 4° to 20° and using the Euler angles assigned to the condensing wing to calculate the reconstructions. A 4° geodesic radius selected 2,375 particles and produced a reconstruction with reasonable modifying and condensing wing but limited overall resolution. Increasing the radius increased particle number and improved the reconstruction of wing, but progressively blurred the modifying wing because of increased angular heterogeneity (Supplementary Fig. 4a, top row). We next applied the e2gmm patch-by-patch refinement algorithm^23^ to test whether heterogeneity within each selected subset could be further resolved. At smaller radii (4°, 8°, and 12°), e2gmm substantially improved overall resolution and recovered clearer density in the modifying wing despite the limited particle numbers. In contrast, at the 20° radius, modifying wing remained poorly resolved after e2gmm refinement, indicating that the selected particles contained relative-orientation heterogeneity beyond what could be corrected by local patch-based refinement alone (Supplementary Fig. 4a, bottom row).

### Landscape of INO80-hexasome complex

Particles from previous reported^19^ three INO80-hexasome classes are combined into a single dataset containing 356,280 particles and processed in RELION 5.0. Appling a soft mask around the INO80 for focused refinement produced an INO80 reconstruction at 2.8 Å. To refine the hexasome, the INO80 signal was subtracted from each particle. Particles after subtraction were recentered to the hexasome and a loose mask covering the hexasome was used for focused refinement, starting from an angular sampling of 15°. This procedure yielded a hexasome-focused reconstruction at 3.1Å. The final STAR files from the INO80-focused and hexasome-focused refinements were imported into cryoROLE. The hexasome was defined as the reference domain and INO80 as the moving domain, and a per-particle RO was computed for each particle. Unless otherwise noted, RO landscapes were displayed with a threshold of 2.0.

The initial INO80-hexasome RO landscape has an elongated shape. Canonicalization in RV space maps the Z axis perpendicular approximately to the surface of hexasome, and Y axis along the direction from Arp5 module to Ino80 ATPase domain. To generate representative reconstructions along the dominant motion coordinate, we sampled the aligned landscape along the primary fixed-axis Euler coordinate α from −60° to +40° in 20° increments. For each target coordinate, particles were selected within a 7–9° geodesic radius, and six reconstructions were calculated from the selected particles. These reconstructions were used to visualize the orientation of INO80 relative to the hexasome.

### Landscape of ribosome

We obtained from authors of two published 70S ribosome datasets^21^: a regular ribosome dataset containing 655,257 particles, and another one of ribosome after temperature annealing containing 509,941 particles. The two datasets were merged for processing in cryoSPARC. The reference map was oriented so that its Z axis coincided with the axis of canonical inter-subunit ratchet motion and its Y axis along the tilt axis defined by ribosome angle decomposition^10^. Because this predefined frame was already aligned with the expected biological motions, no additional RV-axis canonicalization was applied for the ribosome landscapes unless otherwise specified.

A single non-uniform refinement was first performed on the combined particle stack, followed by local refinements using soft masks around the large subunit (LSU), small-subunit body (SSU-body) and small-subunit head (SSU-head), respectively. These refinements yielded maps at 2.28 Å for LSU, 2.44 Å for SSU-body and 2.56 Å for SSU-head for optimal composite map arrangement and provided per-particle orientation estimates for all three rigid bodies.

The *cs*. files were directly input into cryoROLE to calculate the landscape. For each particle, we computed three pairwise ROs: SSU-body relative to LSU (B2L, with LSU as reference), SSU-head relative to SSU-body (H2B, with SSU-body as reference) and SSU-head relative to LSU (H2L, with LSU as reference). RO landscapes were generated from the combined dataset and separately for the unannealed and annealed dataset. All landscapes are displayed with a threshold of SLD = 2.0.

To map published ribosome states onto the RO landscapes, each atomic model was segmented into LSU, SSU-body and SSU-head components using UCSF chimera^24^. In Chimera, the LSU component was first rigidly fitted into the composite map to anchor the model in the common reference frame. From this anchored model, the relative transforms of SSU-body and SSU-head were computed and converted to the same fixed-axis Euler display coordinates. Around each coordinate, particles were selected within a 3° geodesic radius to calculate reconstructions using the .cs of LSU. This procedure enabled direct comparison between the selected cryo-EM reconstructions and the corresponding published structural states. All reconstructions are models were analyzed in ChimeraX^25^.

### CryoDRGN analysis

CryoDRGN analysis was performed on two particle sets from the INO80-hexasome dataset: the full particle set and a landscape-selected subset centered at the RO coordinate (α, β, γ) = (0°, 0°, 0°) with an 8° geodesic radius, containing 16,007 particles. Particle images were down sampled to 128 × 128 pixels before training. For both datasets, cryoDRGN models were trained using a latent dimension of 8 for 25 epochs. Representative volumes were generated from 20 k-means-sampled latent-space positions.

## Data Availability

Particle stack of hFASN and INO80-Hexasome complex used in this study have been deposited to EMPIAR with accession code of xxx (hFASN) and 13299 (INO80-hexasome).

## Code availability

The program CryoROLE (cryo-EM relative orientation landscape) used in this study is open source and available via GitHub at https://github.com/yifancheng-ucsf/cryorole with instructions.

## ACKNOWLEDGMENTS

We thank Dr. Q. Shen for providing the ribosome datasets for our test. This work was supported by grants from the National Institute of Health (R35GM140847, and U54AI170792) to Y.C. The UCSF cryo-EM facility was partially supported by NIH grants (S10OD020054, S10OD021741, and S10OD025881). Y.C. is an investigator of the Howard Hughes Medical Institute.

## AUTHOR CONTRIBUTIONS

C.L., W.C. and Y.C. conceptualized project and designed experiments. C.L. developed cryoROLE program, with suggestions and helps from W.C., H.W. and Y.C. C.L. and W.C. worked together on FASN. C.L. and H.W. worked together on INO80-hexasome complex. All authors wrote the manuscript.

## Competing interests

Y. Cheng is a non-shareholder member of Scientific Advisory Board at ShuiMu BioSciences Ltd. and Pamplona Therapeutic Co. Ltd. All other authors declare no competing interest.

**Correspondence and requests for materials** should be addressed to Yifan Cheng.

---

**Supplementary Figure 1.**
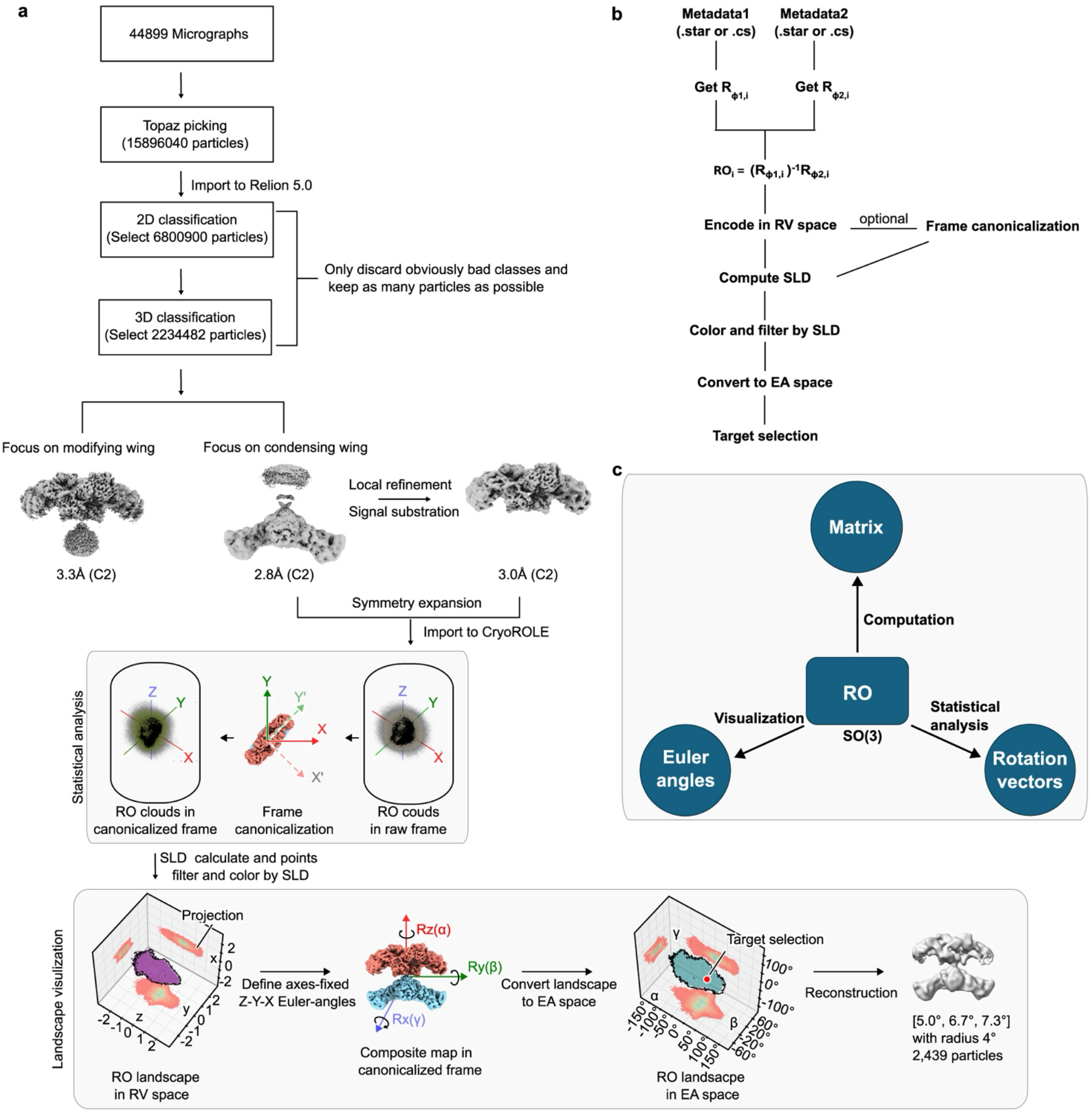
Human FASN processing and cryoROLE workflow. **a**, Workflow of processing FASN dataset and generation of RO landscape by cryoROLE. **b**, cryoROLE computational pipeline. **c**, RO representations used for computation, statistical analysis and visualization.

**Supplementary Figure 2.**
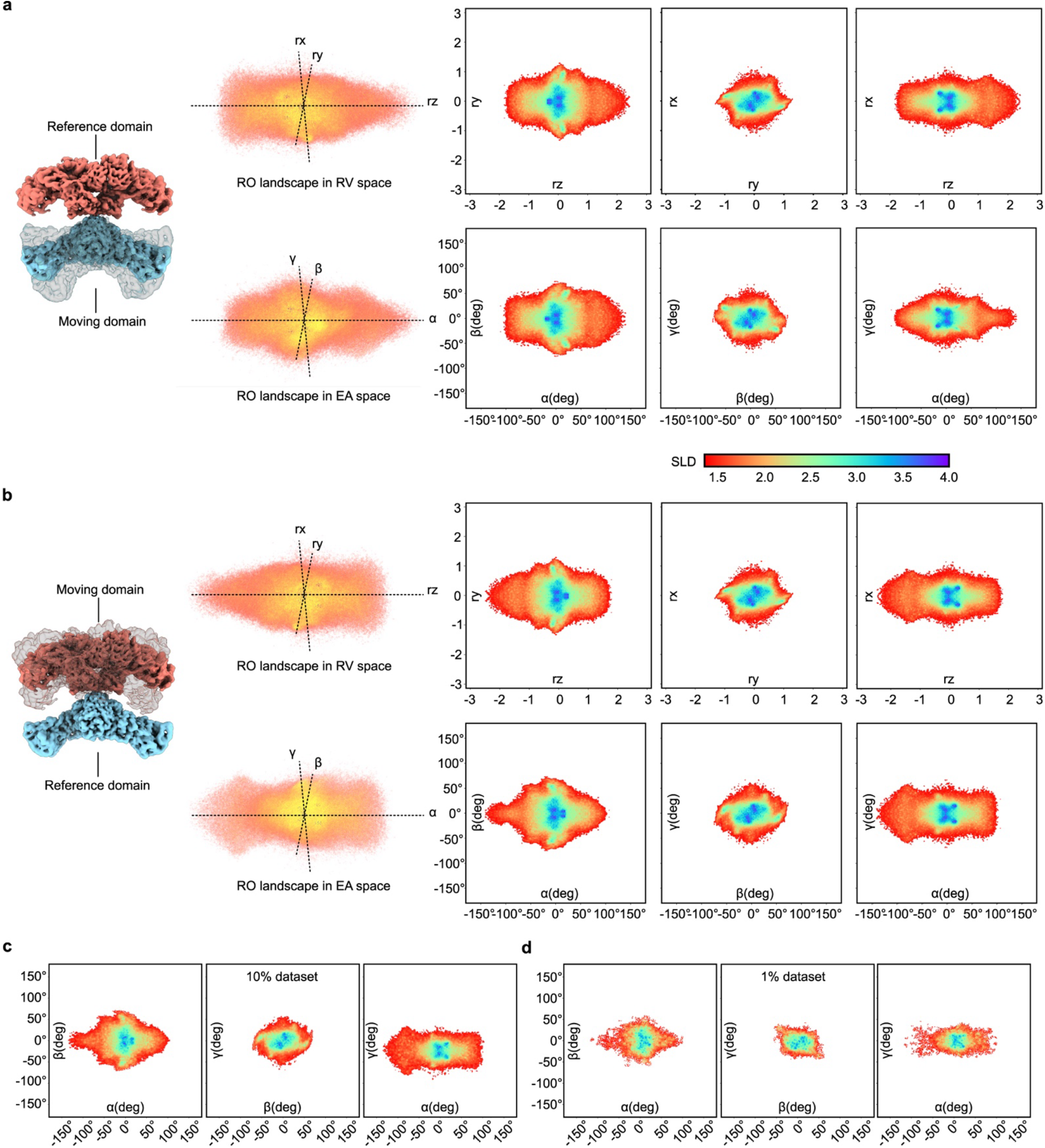
Effect of exchanging reference and moving domains. **a and b**, Moving and reference domains are swapped in the composite map of hFASN, producing an inverted landscape in RV space and the corresponding re-parameterized landscape in EA space, without changing the intrinsic landscape structure. **c and d**, RO landscapes from random 10% and 1% subsets of the FASN dataset.

**Supplementary Figure 3.**
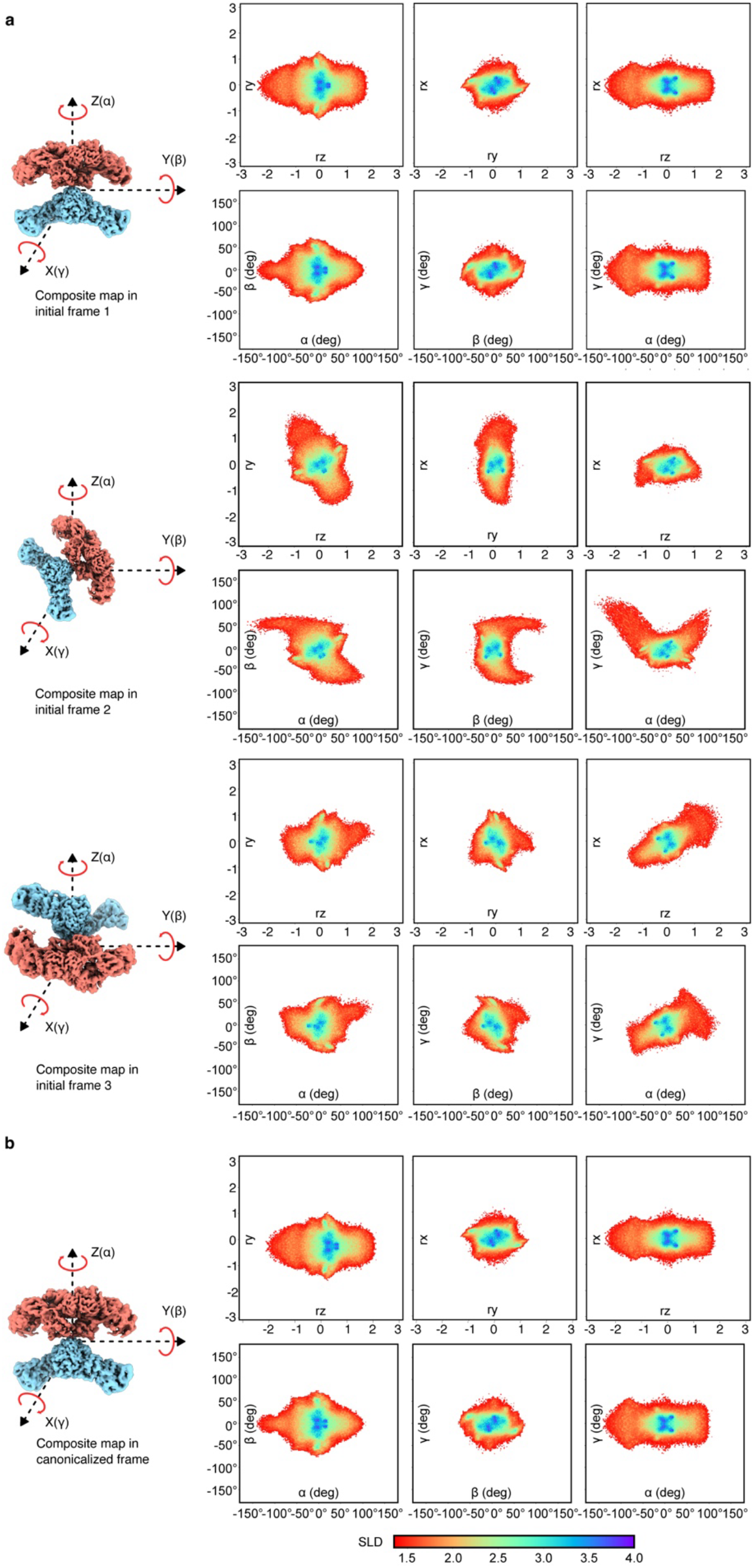
Effect of reorienting the reference composite map. **a**, RO landscapes of the same FASN dataset displayed using three different initial composite-map coordinate frames. Changing the initial frame rotates the RO cloud linearly in rotation-vector (RV) space and produces nonlinear changes in Euler-angle displays. **b**, After canonicalization, the corresponding RO landscapes converge to the same canonical representation.

**Supplementary Figure 4.**
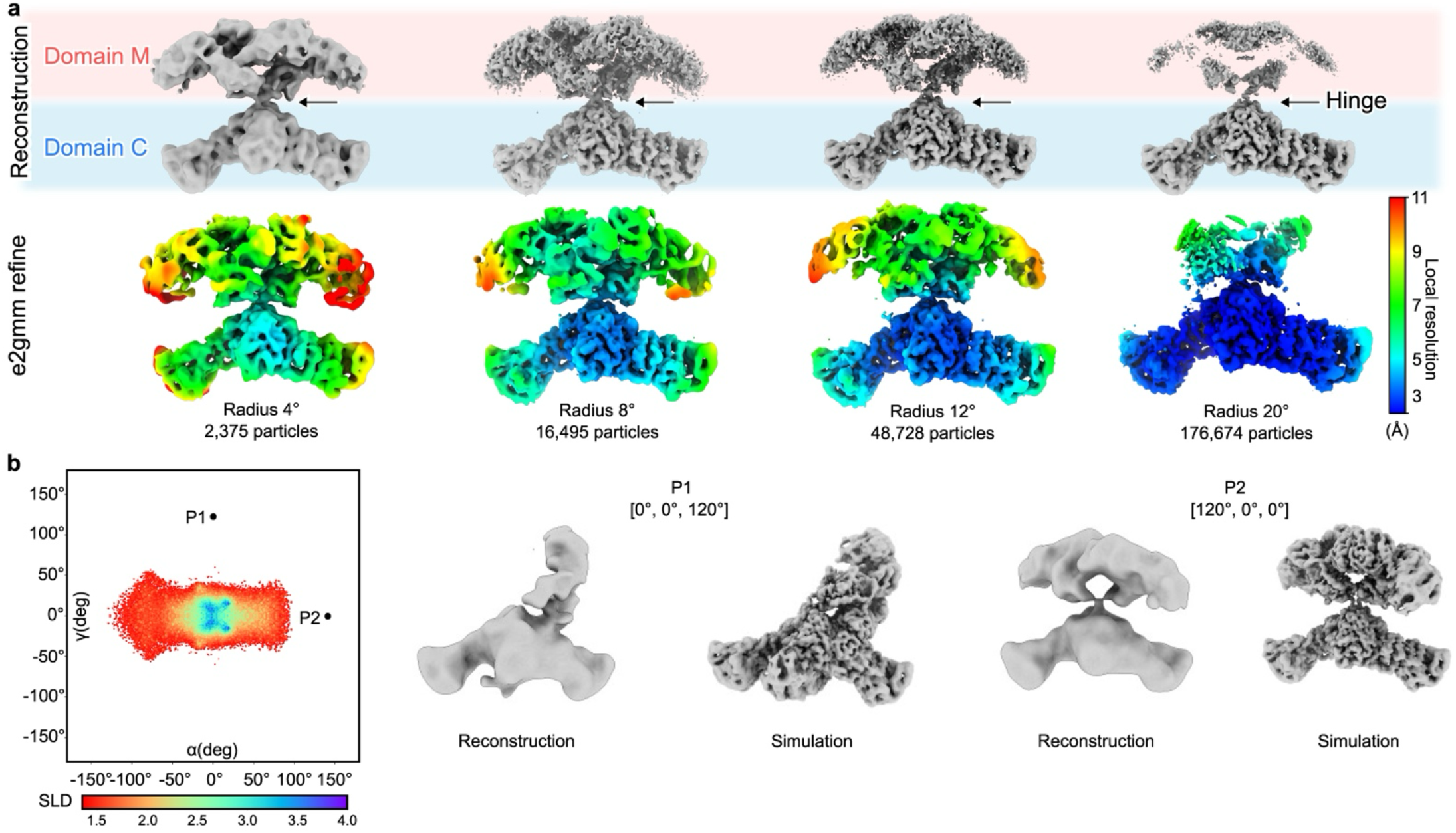
Practical considerations for RO landscape-guided reconstruction. **a**, Reconstructions from particles selected around the same FASN RO coordinate using increasing geodesic radii. Larger radii increase particle number but reduce RO homogeneity, leading to progressive blurring of the moving domain. Bottom row, e2GMM-refined maps colored by local resolution. **b**, Reconstructions from low-SLD coordinates compared with corresponding simulations.

**Supplementary Figure 5.**
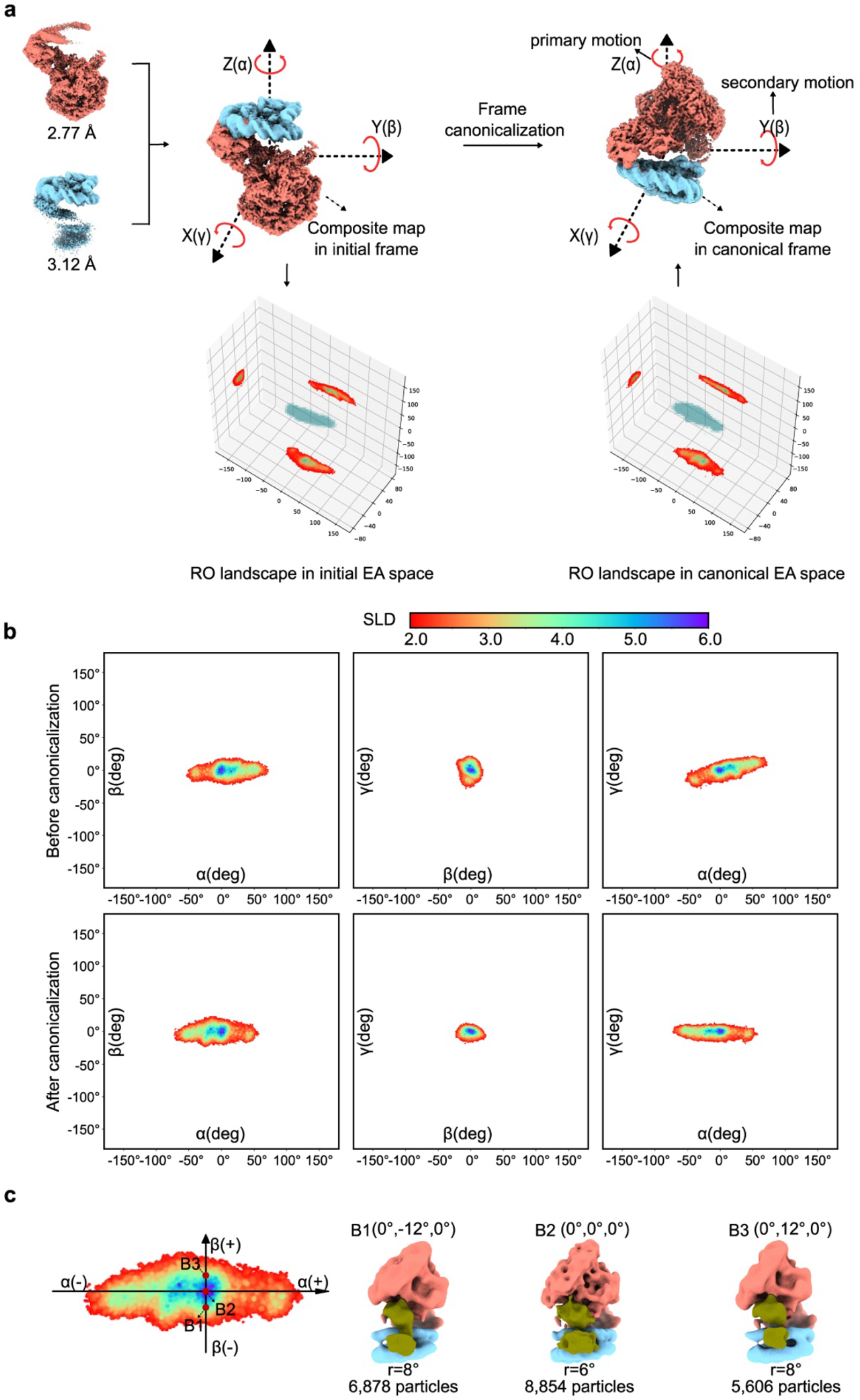
Motion-aligned frame canonicalization of the INO80-hexasome RO landscape. **a**, INO80 and hexasome focused-refinement maps were assembled into a composite map and reoriented from the initial frame to the canonical frame. Corresponding RO landscapes are shown in fixed-axis Z–Y–X Euler-angle space. **b**, Two-dimensional RO projections before and after canonicalization. **c**, Reconstructions from particles selected along the secondary β coordinate at B1, B2 and B3.

**Supplementary Fig. 6.**
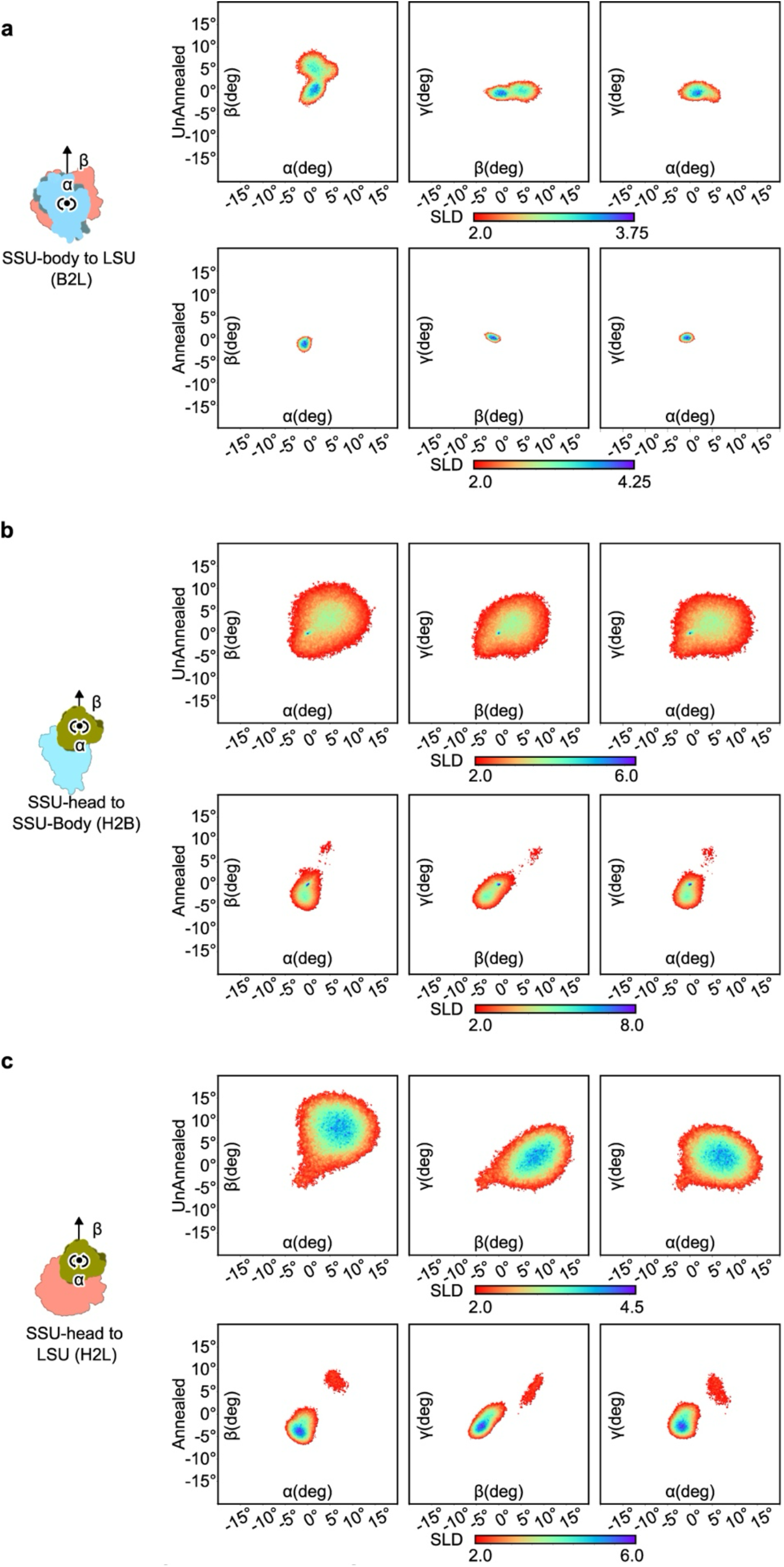
RO landscapes of ribosomal domain motions without and after temperature annealing. **a–c**, Two-dimensional projections of ribosome RO landscapes for the unannealed and annealed datasets. Three pairwise domain motions are SSU-body to LSU (B2L, **a**), SSU-head to SSU-body (H2B **b**,) and SSU-head to LSU (H2L, **c**).

**Supplementary Fig. 7.**
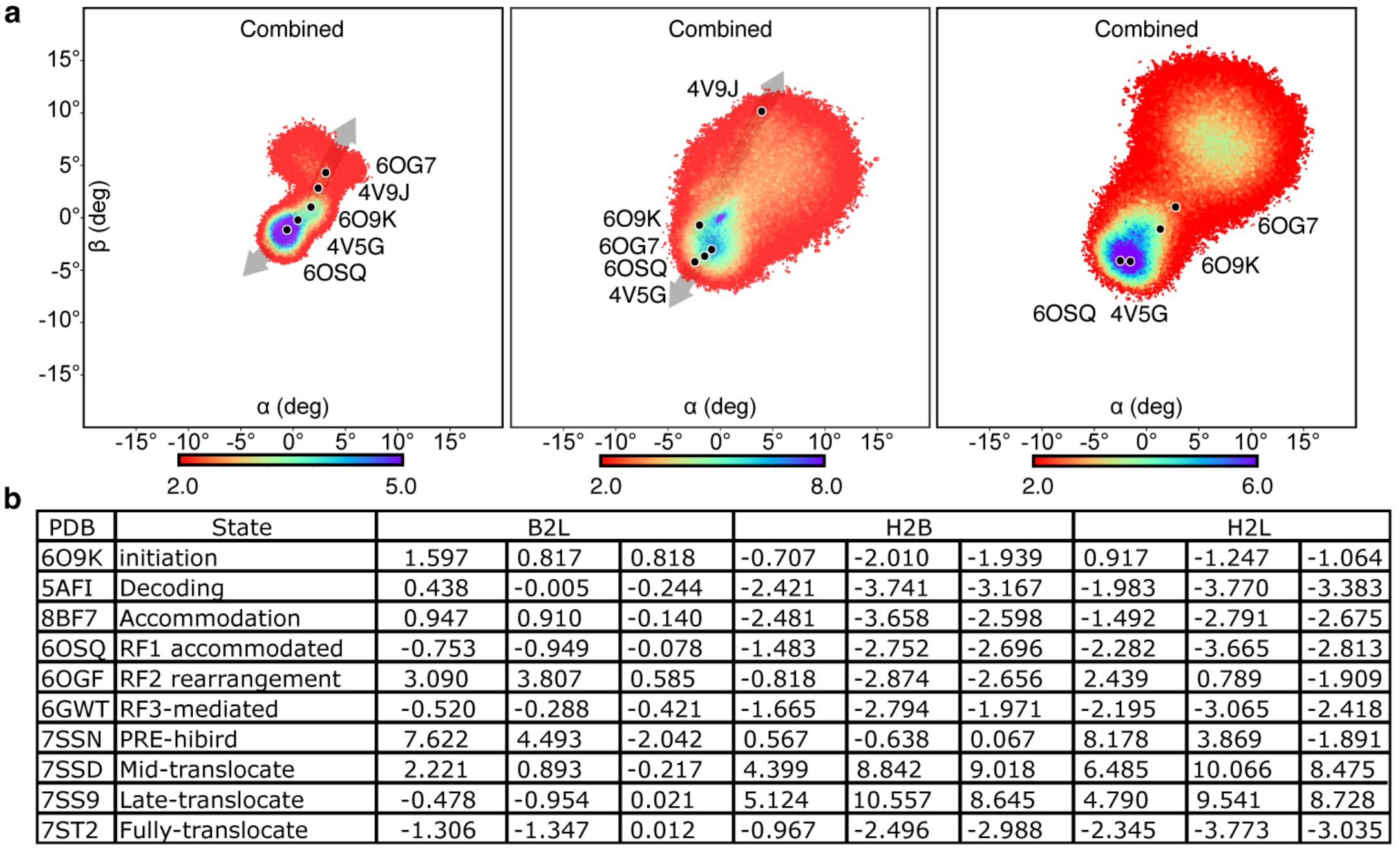
Mapping known ribosome states onto the combined RO landscapes. **a**, Published *E. coli* 70S ribosome structures were mapped onto the combined RO landscapes for B2L, H2B, and H2L motions. Black dots mark the mapped coordinates of representative functional states. **b**, Table of mapped α, β, and γ coordinates for each PDB structure in the three RO landscapes.

**Supplementary Fig. 8.**
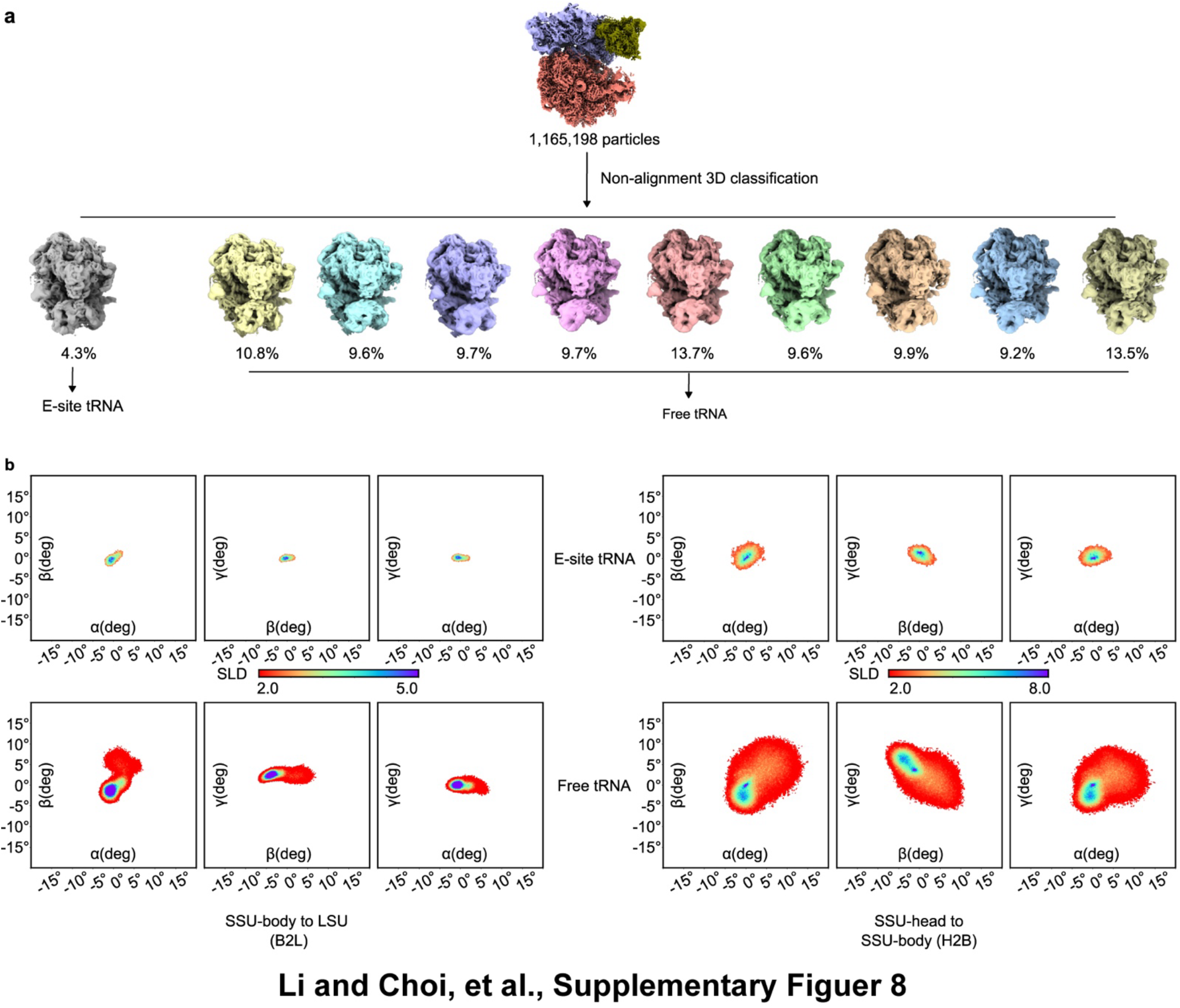
E-site tRNA binding is associated with a restricted ribosome RO landscape. **a**, The combined ribosome dataset was subjected to non-alignment 3D classification. One minor class (4.3%) showed clear density for an E-site tRNA, whereas the remaining classes lacked detectable tRNA density and were grouped as the tRNA-free population. **b**, Two-dimensional projections of the RO landscapes of B2L and H2B for the E-site tRNA-bound and tRNA-free populations.

**Supplementary Fig. 9.**
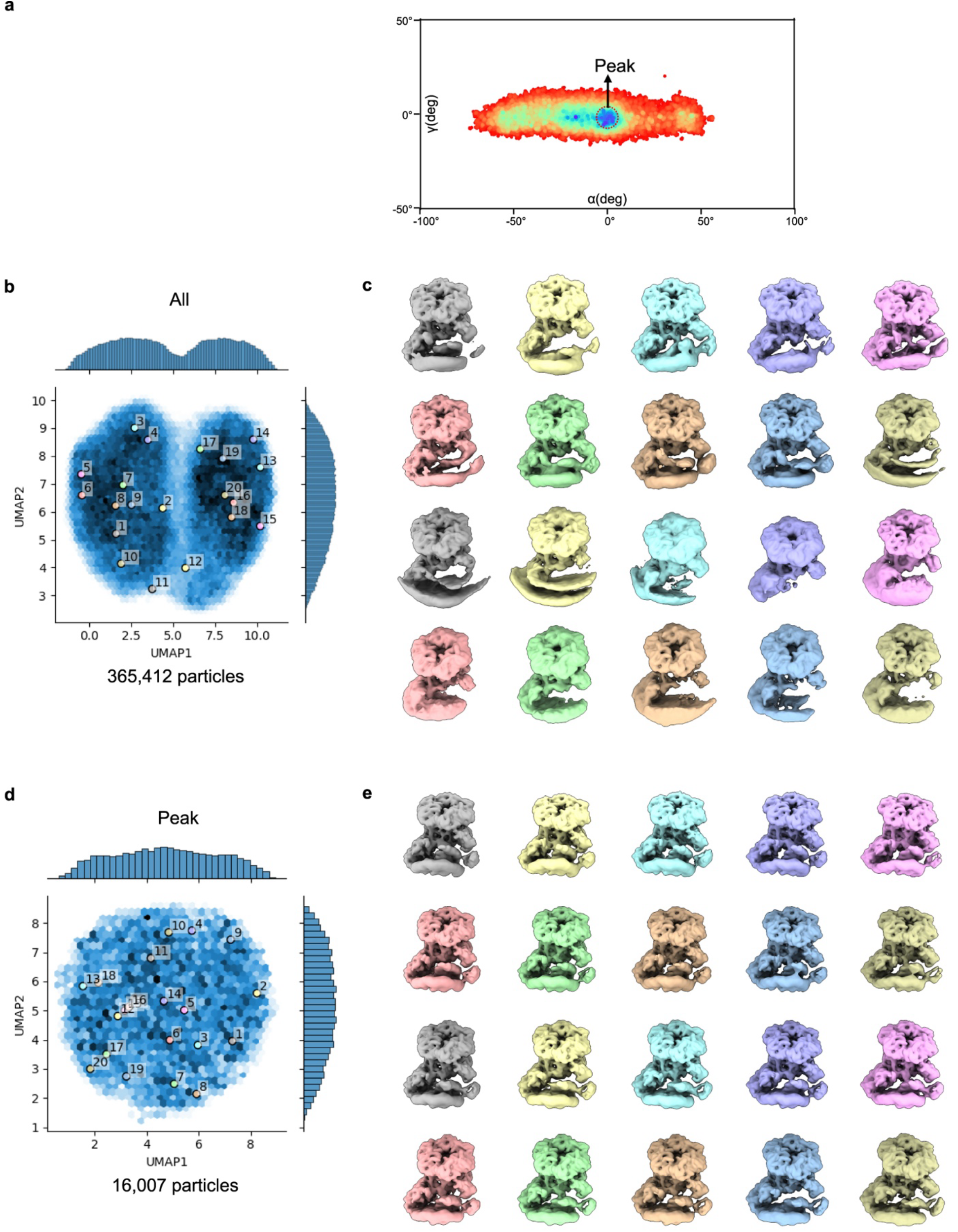
CryoDRGN analysis of the full INO80-hexasome dataset and of a cryoROLE-selected local subset. **a**, Peak (0°,0°,0°) in the canonicalized RO landscape. **b** and **c**, UMAP embedding and k-means (k=20) reconstructions from cryoDRGN analysis of the full dataset (365,412 particles). **d** and **e**, UMAP embedding and k-means ((k=20)) reconstructions from cryoDRGN analysis of particles selected within an 8° geodesic radius around Peak (16,007 particles). Both analyses were performed with the same default cryoDRGN parameters.

